# A Phylogenomic Framework for Charting the Diversity and Evolution of Giant Viruses

**DOI:** 10.1101/2021.05.05.442809

**Authors:** Frank O. Aylward, Mohammad Moniruzzaman, Anh D. Ha, Eugene V. Koonin

## Abstract

Large DNA viruses of the phylum *Nucleocytoviricota* have recently emerged as important members of ecosystems around the globe that challenge traditional views of viral complexity. Numerous members of this phylum that cannot be classified within established families have recently been reported, and there is presently a strong need for a robust phylogenomic and taxonomic framework for these viruses. Here we report a comprehensive phylogenomic analysis of the *Nucleocytoviricota*, present a set of giant virus orthologous groups (GVOGs) together with a benchmarked reference phylogeny, and delineate a hierarchical taxonomy within this phylum. We show that the majority of *Nucleocytoviricota* diversity can be partitioned into 6 orders, 32 families, and 344 genera, substantially expanding the number of currently recognized taxonomic ranks for these viruses. We integrate our results within a megataxonomy that has been adopted for all viruses to establish a unifying framework for the study of *Nucleocytoviricota* diversity, evolution, and environmental distribution.

## Main Text

Large double-stranded DNA viruses of the phylum *Nucleocytoviricota* are a diverse group of viruses with virion sizes reaching up to 1.5 μm and genome sizes up to 2.5 Mb, comparable to many bacteria and archaea as well as picoeukaryotes [1–4]. This phylum currently includes two classes that encompasses five orders with seven recognized and several additional, proposed families. The viruses in the families *Poxviridae, Asfarviridae*, and *Iridoviridae* infect metazoans, whereas those in the families *Mimiviridae, Phycodnaviridae* and Marseilleviridae as well as several putative families primarily infect algae or heterotrophic unicellular eukaryotes [5–7]. The members of the *Nucleocytoviricota* span an exceptionally broad range of genome sizes from below 100 kb to more than 2.5 Mb. Many comparative genome analyses have documented the highly complex, chimeric nature of their genomes whereby numerous genes appear to have been acquired from diverse cellular lineages and other viruses [8–12]. These multiple, dynamic gene exchanges between viruses and their hosts [13–16] as well as the large phylogenetic distances between different families and especially orders [11,17,18] make the investigation of the evolution and taxonomic classification of the *Nucleocytoviricota* a challenging task. Despite these difficulties, early comparative genomic analyses studies succeeded in identifying a small set of core genes that could be reliably used to produce phylogenies that encompass the entire diversity of *Nucleocytoviricota*, leading to the conclusion that all these viruses share common evolutionary origins [17,19].

Recent studies have reported numerous new *Nucleocytoviricota* genomes, many of which seem to represent novel lineages with only distant phylogenetic affinity for previously identified taxa [9,15,20]. For example, many viruses that infect a variety of protist genera have been discovered that bear phylogenetic affinity to *Mimiviridae*, but do not fall within the same clade as the canonical Acanthamoeba polyphaga Mimivirus [8,21,22]. Moreover, numerous metagenome-assembled genomes (MAGs) have been reported that also appear to form novel sister clades to the *Mimiviridae, Asfarviridae* and other families [9,15,20]. The uncertainty of the phylogenetic structure and taxonomy *Nucleocytoviricota* is a major impediment for the ongoing efforts to characterize the diversity of these viruses in the environment, as well as studies that seek to better understand the evolutionary origins of unique traits within this viral phylum. As more studies begin to chart the environmental diversity of *Nucleocytoviricota*, defining taxonomic groupings that encompass equivalent phylogenetic breadths will be critical for the exploration of the geographic and temporal variability in viral diversity, and for comparing results from different studies. Moreover, the evolutionary origins of large genomes, virion sizes, and complex metabolic repertoires in many *Nucleocytoviricota* are of great interest, and ancestral state reconstructions as well as tracking horizontal gene transfers fully depend on a robust phylogenetic framework.

Here we present a phylogenomic framework for charting the diversity and evolution of *Nucleocytoviricota*. We first assess the strength of the phylogenetic signals from different marker genes that are found in a broad array of distantly related viruses, and arrive at a set of 7 genes that performs well in our benchmarking of concatenated protein alignments. Using this hallmark gene set, we then perform a large-scale phylogenetic analysis and clade delineation of the *Nucleocytoviricota* to produce an hierarchical taxonomy. Our taxonomy includes the established families *Poxviridae, Asfarviridae, Iridoviridae, Phycodnaviridae, Marseilleviridae*, and *Mimiviridae* as well as 26 proposed new family-level clades and one proposed new order. Sixteen of the families are represented only by genomes derived from cultivation- independent approaches, underscoring the enormous diversity of these viruses in the environment. We integrate these family-level classifications into the broader megataxonomy framework that has been adopted for all viruses [3] to arrive at a unified and hierarchical classification scheme for the entire phylum *Nucleocytoviricota*.

## Results

### Phylogenetic Benchmarking of Marker Genes

We first generated a dataset of protein families to identify phylogenetic marker genes that are broadly represented across *Nucleocytoviricota*. To this end, we selected a set of 1,380 quality-checked *Nucleocytoviricota* genomes that encompassed all of the established families (Table S1, see Methods). By clustering the protein sequences encoded in these genomes (see Methods for details), we then generated a set of 8,863 protein families, which we refer to as Giant Virus Orthologous Groups (GVOGs). We examined 25 GVOGs that were represented in >70% of all genomes and ultimately arrived at a set of 9 GVOGs that were potentially useful for phylogenetic analysis, which is largely consistent with the previous studies that have identified phylogenetic marker genes in *Nucleocytoviricota* [18,19,23] (Table 1, Figs S1-S25, see Methods for details; descriptions of all GVOGs provided in Table S1). These GVOGs included 5 genes that we have previously used for phylogenetic analysis of *Nucleocytoviricota*: the family B DNA Polymerase (PolB), A32-like packaging ATPase (A32), virus late transcription factor 3 (VLTF3), superfamily II helicase (SFII), and major capsid protein (MCP) [9]. In addition, this set included the large and small RNA polymerase subunits (RNAPL and RNAPS, respectively), the TFIIB transcriptional factor (TFIIB), and the Topoisomerase family II (TopoII).

We evaluated individual marker genes and concatenated marker sets using the Internode Certainty and Tree Certainty metrics (IC and TC, respectively), which provide a measure of the phylogenetic strength of each individual marker gene and have been shown to yield more robust estimates of confidence than the traditional bootstrap [24,25]. The TC values were highest for the RNAP subunits, PolB, and TopoII (Fig 1a), consistent with the view that, in most cases, longer genes carry a stronger phylogenetic signal, apparently thanks to the larger number of phylogenetically-informative characters. The MCP marker had markedly lower TC values than PolB, TopoII, or either of the RNAP subunits; this is likely to be the case because *Nucleocytoviricota* genomes often encode multiple copies of MCP, which complicates efforts to distinguish orthologs from paralogs (Fig. 1a). SFII, TFIIB, A32, and VLTF3 showed lower TC values than the other 5 markers, but these were also the shortest marker genes and would not be expected to yield high quality phylogenies when used individually.

**Figure 1.**
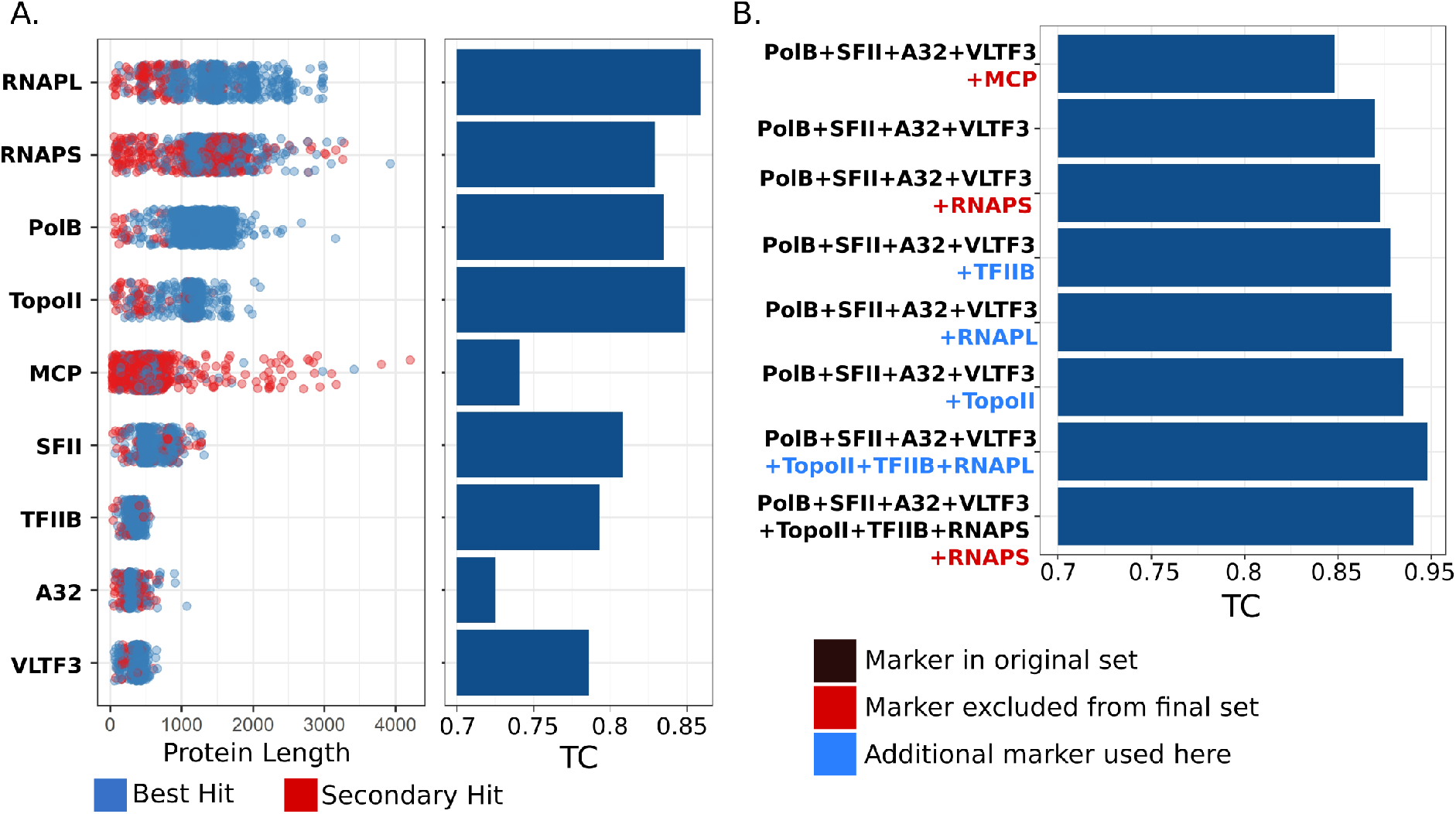
Benchmarking of phylogenetic marker genes for the *Nucleocytoviricota*. A) Dotplot of protein lengths for each of the 9 marker genes examined. Blue dots represent proteins that were the best hit against marker gene HMMs and likely represent true orthologs, while red dots represent multiple copies of marker genes present in a genome. The Tree Certainty (TC) scores of the markers is presented on the barplot on the right. B) TC values for phylogenies made from concatenated alignments of different marker sets. Black text denotes markers we have used previously, red text denotes markers that we not included in the final set, and blue text denotes additional markers used here compared to our original 5-gene set. Note that MCP was used in our original marker set but is excluded from the final 7-gene set.

Next we sought to identify which marker genes provide for the best phylogenetic inference when used in a concatenated alignment. If markers produce incongruent phylogenetic signals, they will yield trees with low TC values when concatenated, even if the individual phylogenetic strength of the markers is high [25]. We started by assessing the TC of the 5-gene set that we have previously used. Surprisingly, the TC of this set was lower than that of some individual markers (TC of 0.865; Fig. 1b), suggesting that some of the markers provide incongruent signals. We surmised that this was most likely due to the MCP, given the large number of paralogs of this protein found in some *Nucleocytoviricota* (Fig 1a). As we suspected, removal of MCP increased the TC of the concatenated tree (from 0.865 to 0.875), indicating that this gene should be excluded from concatenated alignments (Fig 1b). The addition of the RNAPS, RNAPL, TFIIB, or TopoII markers to the 4-gene set increased the TC (Fig 1b) although a 7-gene marker set that excluded RNAPS performed best overall (TC of 0.898). This 7-gene marker set therefore represents a substantial improvement over the initial 5-gene set, and we used these genes for subsequent phylogenetic analysis and clade demarcation.

### An Hierarchical Taxonomy for *Nucleocytoviricota*

The best-quality phylogenetic tree produced with the 7-gene marker set could be broadly divided into two class-level and 6 order-level clades, five of which were consistent with the orders in the recently adopted megataxonomy of viruses (Fig. 2) [3]. The *Chitovirales* and *Asfuvirales* orders, which respectively contain the *Poxviridae* and *Asfarviridae*, formed a distinct group with a long stem branch (class *Pokkesviricetes*) that we used to root the tree, consistent with previous studies [19,26]. The *Pimascovirales*, which includes Pithoviruses, Marseilleviruses and *Iridoviridae/Ascoviridae*, also formed a highly-supported monophyletic group. The current order *Algavirales*, which includes the *Phycodnaviridae*, Chloroviruses, Pandoraviruses, Molliviruses, Prasinoviruses, and Coccolithoviruses, was paraphyletic, and we split this order into two groups based on their placement in the phylogeny. In the proposed taxonomy, we retain the existing *Algavirales* name for the clade that contains the Chloroviruses and Prasinoviruses, and additionally propose the order *Pandoravirales* for the group that includes the Pandoraviruses and Coccolithoviruses. The *Imitervirales*, which contain the *Mimiviridae*, formed a sister group to the *Algavirales*.

**Figure 2.**
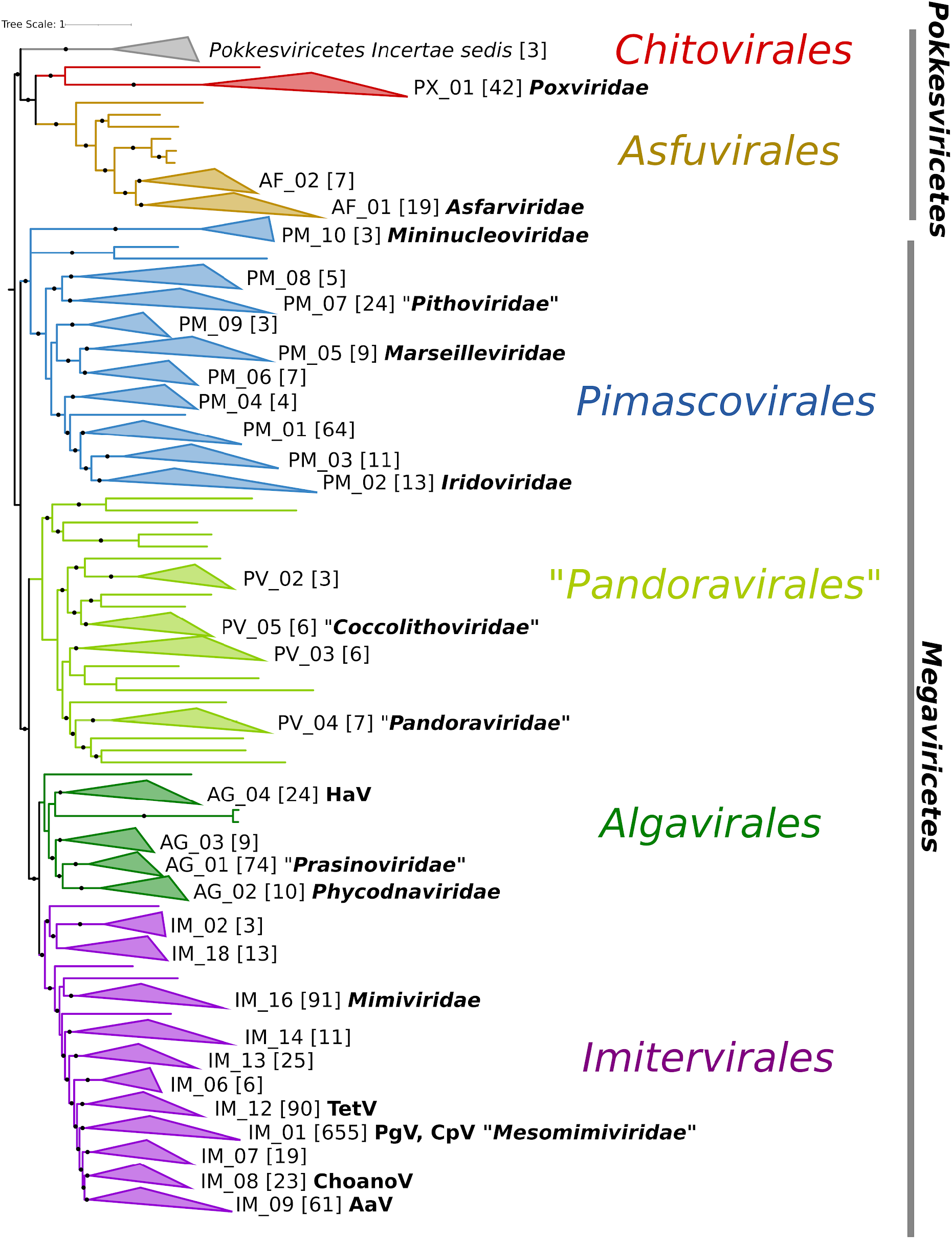
Phylogeny of *Nucleocytoviricota* based on the 7-gene marker gene set that had the highest TC value of those tested. The phylogeny was inferred using the LG+I+F+G4 model in IQ-TREE. Solid circles denote IC values >0.5. Families are denoted by collapsed clades, with their non-redundant identifier provided at their right. The number of genomes in each clade is provided in brackets. Established family names are provided in bold italics, and family names proposed here are provided in quotes. The presence of notable cultivated viruses are provided in bold next to some clades. Abbreviations: HaV: Heterosigma akashiwo virus, PgV: Phaeocystis globosa virus, CpV: Chrysochromulina parva virus, TetV: Tetraselmis virus, ChoanoV: Choanoflaggelate virus, Aav: Aureococcus anophagefferens virus.

From our reference tree we delineated taxonomic levels using the Relative Evolutionary Distance (RED) of each clade as a guide, using an approach similar to the one recently employed for bacteria and archaea [27]. RED values vary between 0 and 1, with lower values denoting phylogenetically broad groups that branch closer to the root and higher values denoting phylogenetically shallow groups that branch closer to the leaves. The RED of the *Nucleocytoviricota* classes ranges from 0.017-0.032, whereas the values for the orders range from 0.158-0.240 (Fig. 3a). We delineated family- and genus-level clades so that they had non-overlapping RED values that were higher than their next-highest taxonomic rank (Fig 3a). This approach yielded clades that were consistent with families and genera currently recognized by the International Committee on Taxonomy of Viruses (ICTV [28]), such as the *Chlorovirus, Prasinovirus*, and *Mimiviridae* (see below; full classification information in Table S2). To ensure that putative families were not defined by spurious placement of individual genomes, we accepted only groups with ≥ 3 members and left other genomes in the tree as singletons with *incertae sedis* as the family identifier. This approach yielded a total of 32 families, not including 22 singleton genomes that potentially represent additional families and are listed as *incertae sedis* here. Moreover, at the genus-level this yielded 344 genera, including 213 genera with a single representative (Fig 2, Fig 3).

**Figure 3.**
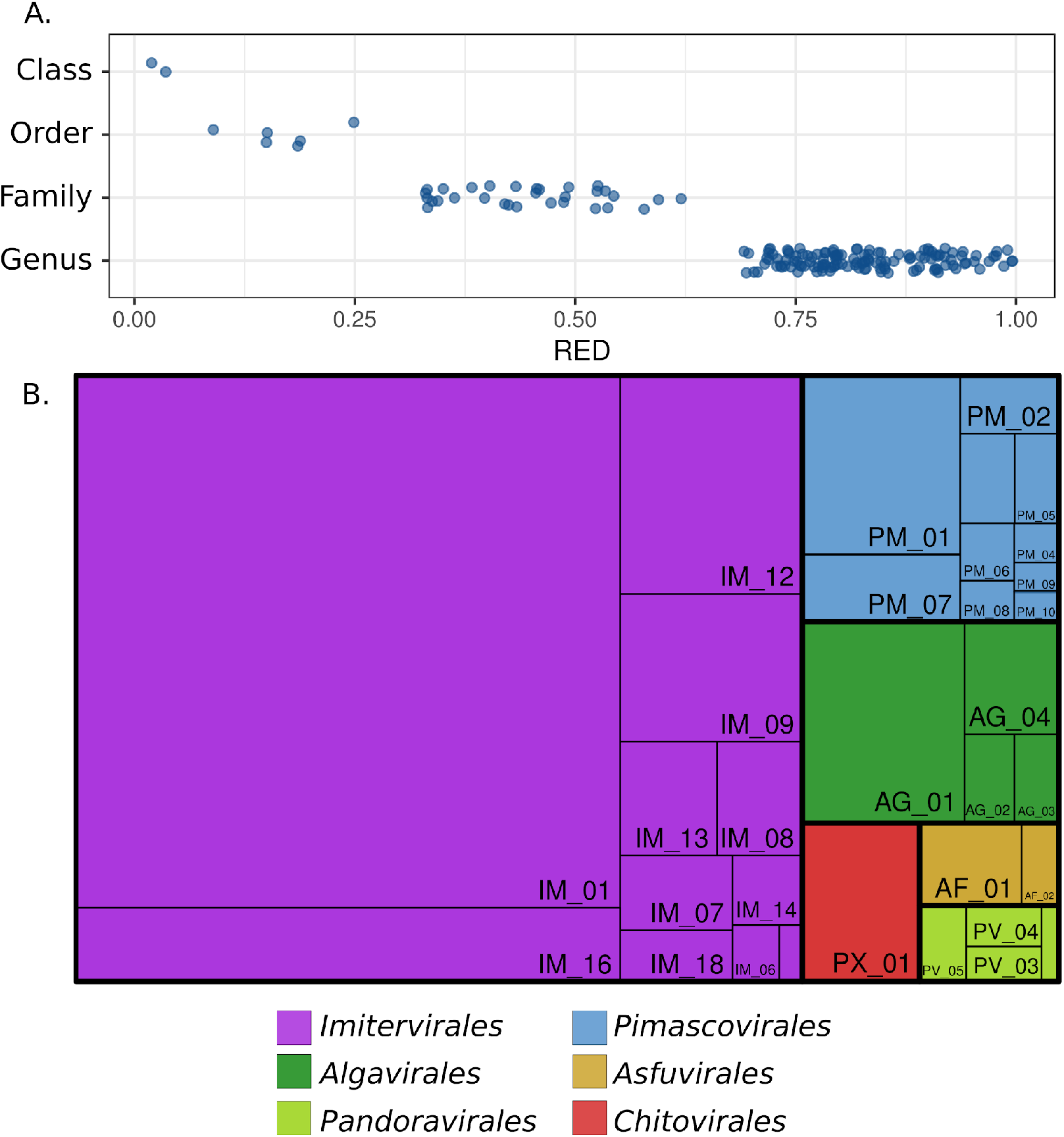
Summary of the *Nucleocytoviricota* taxonomy. A) Relative Evolutionary Divergence (RED) values for *Nucleocytoviricota* classes, orders, and families. B) Treemap diagram of the *Nucleocytoviricota* in which orders and families are shown. The area of each rectangle is proportional to the number of genomes in the respective taxon.

Of the 32 families, 6 correspond to the families currently recognized by the ICTV, for which we retained the existing nomenclature (*Asfarviridae, Poxviridae, Marseilleviridae, Iridoviridae, Phycodnaviridae*, and *Mimiviridae*). The *Ascoviridae* are included within the *Iridoviridae*, and so we use the latter family name here. In addition, we propose six family names here: “Prasinoviridae”, which include the prasinoviruses, “Pandoraviridae”, which include the Pandoraviruses and Mollivirus sibericum, “Coccolithoviridae”, which include the coccolithoviruses, “Pithoviridae”, which include Pithoviruses, Cedratviruses, and Orpheoviruses, “Mesomimiviridae”, which includes several haptophyte viruses previously defined as “extended *Mimiviridae*”, and “Mininucleoviridae”, which includes several viruses of Crustacea [22]. Some of these family names have been used previously, such as *Pandoraviridae* and *Mininucleoviridae*, but so far have not been formally recognized by the ICTV. For other proposed families, we provide non-redundant identifiers corresponding to their order, and we anticipate future studies will provide information for selecting appropriate family names once more is learned on the host ranges and molecular traits of these viruses. Two of the families contained only a single cultivated representative (AG_04 and IM_09), whereas 16 families included none.

Notably, the *Imitervirales* contain 11 families, as well as 4 singleton viruses that likely represent additional family-level clades. This underscores the vast diversity of the large viruses in this group, which is consistent with the results of several studies reporting an enormous diversity of *Mimiviridae*-like viruses in the biosphere, in particular in aquatic environments [9,29–31]. Other studies have suggested additional nomenclature to refer to these *Mimiviridae*-like viruses, such as the ‘extended *Mimiviridae’* and the sub-families *Mesomimivirinae*, or *Megamimivirinae*, but our results suggest that an extensive array of new families is warranted within *Imitervirales*, given the broad genomic and phylogenetic diversity within this group. Several of the proposed new families contain representatives that have recently been described; IM_12 contains the Tetraselmis virus (TetV), which encodes several fermentation genes [10], IM_09 contains Aureococcus anophagefferens virus (AaV), which is thought to play an important role in brown tide termination [32], and IM_08 contains a virus of Choanoflagellates [33] (Fig 2). Family IM_01 contains cultivated viruses that infect haptophytes of the genera *Chrysochromulina* and *Phaeocystis*, which were previously proposed to be classified in the subfamily *Mesomimivirinae* [22]. We propose the name *Mesomimiviridae* to denote the family-level status of this lineage, while still retaining reference to this original name. Notably, the *Mesomimiviridae* includes by far the largest total number of genomic representatives in our analysis (655, including 652 MAGs; Fig 2, Fig 3b), the vast majority of which are derived from aquatic environments (Fig S26), suggesting that members of this family are important components of global freshwater and marine ecosystems.

All families within the *Imitervirales* except one included members with genome sizes >500 kbp, highlighting the giant genomes that are characteristic of this lineage (Fig 4a). Genes involved in translation, including tRNA synthetases and translation initiation factors, were consistently highly represented in the *Imitervirales*, showing that the rich complement of these genes that has been described for the *Mimiviridae* is broadly characteristic of other families in this order (Fig 4b) [34,35]. Genes involved in glycolysis and the TCA cycle, cytoskeleton components, such as viral-encoded actin, myosin, and kinesin proteins, and nutrient transporters including those that target ammonia and phosphate were also common in the *Imitervirales* (Fig 4b), underscoring the complex functional repertoires of this virus order.

**Figure 4.**
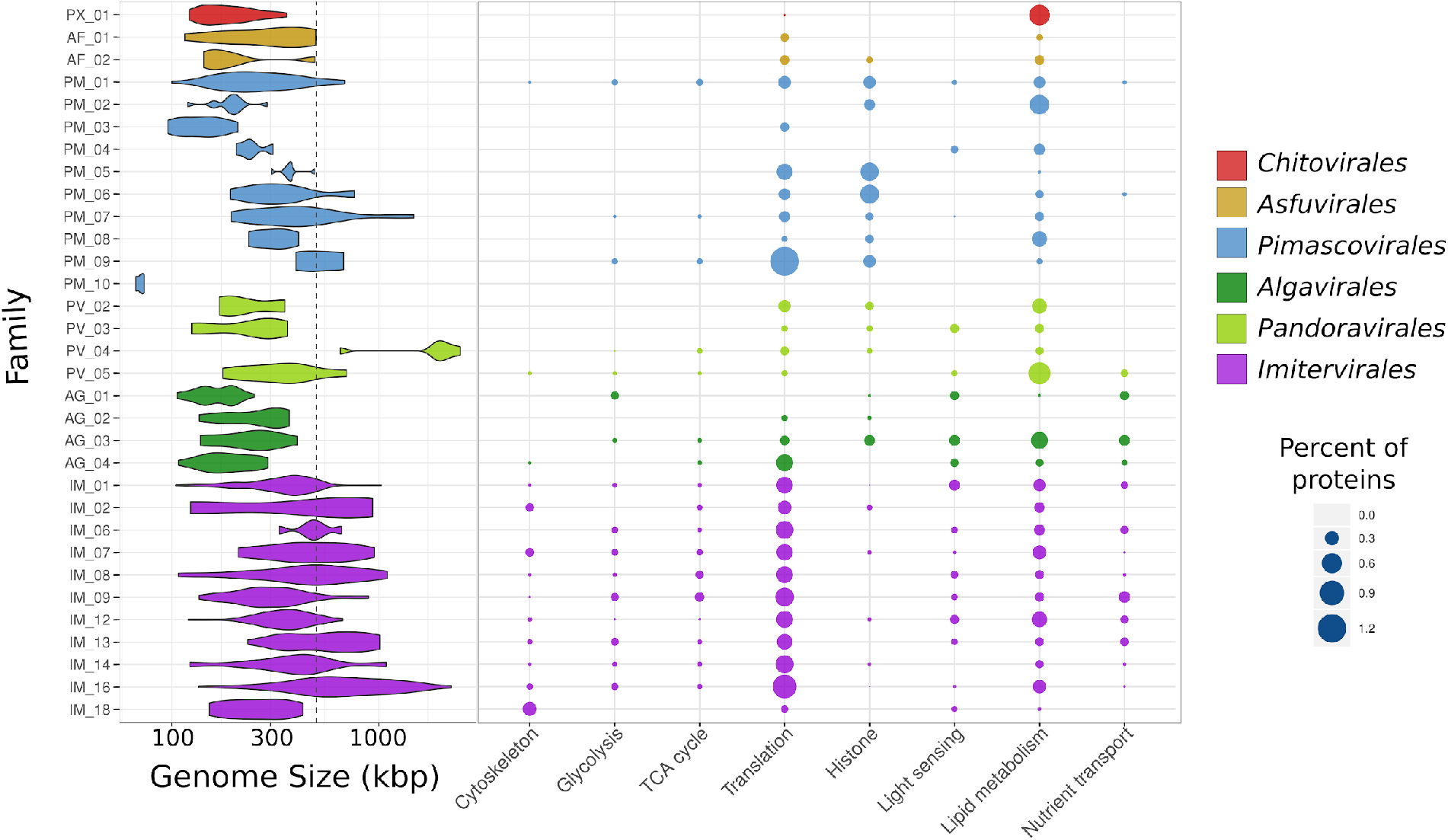
Genomic characteristics of the *Nucleocytoviricota*. A) Violin plot showing the genome size distribution across the Nucleocytoviricota families. The dashed grey line denotes 500 kbp. B) Bubble plot showing the percent of total proteins in each family that could be assigned to GVOGs that belonged to particular functional categories (details in Table S3).

The *Algavirales* is a sister lineage to the *Imitervirales* that contains 4 families encompassing several well-studied algal viruses. The *Prasinoviridae* (AG_01) is a family that includes viruses known to infect the prasinophyte genera *Bathycoccus, Micromonas*, and *Ostreococcus [7]*, and cultivation-independent surveys have provided evidence that the MAGs in this clade are also associated with prasinophytes [36]. Similarly, our approach yielded a well-defined *Phycodnaviridae* family (AG_02) composed mostly of chloroviruses, consistent with the similar host range of these viruses [37]. All four families of the *Algavirales* have smaller genome sizes compared to the *Imitervirales* (Fig 4a), but there were still several similarities in their encoded functional repertoires. As noted previously [9,16,33], genes involved in light sensing, including rhodopsins and chlorophyll-binding proteins, were common across the *Imitervirales* and *Algavirales*, perhaps because many of the viruses are found in sunlit aquatic environments where manipulation of host light sensing during infection is advantageous. Moreover, genes involved in nutrient transport, translation, and even some components of glycolysis and the TCA cycle were found in the *Algavirales*, consistent with the complex repertoires of metabolic genes that have been reported for some of these viruses despite their relatively small genome sizes [38,39].

The *Pandoravirales*, a new order we propose here, consists of 4 families, including the *Pandoraviridae* and the *Coccolithoviridae*. The *Pandoraviridae* (PV_04) include *Mollivirus sibericum* as well as the Pandoraviruses, which possess the largest viral genomes known [40]. Grouping of these viruses together in the same family is consistent with previous studies that have shown that *M. sibericum* and the Pandoraviruses have shared ancestry [41,42], and comparative genomic analysis that have shown that they all encode a unique duplication in the glycosyl hydrolase that has been co-opted as a major virion protein in the Pandoraviruses [43]. The *Coccolithoviridae* (PV_05) is mostly comprised of viruses that infect the marine coccolithophore *Emiliania huxleyi*; although much smaller than the genomes of the Pandoraviruses, genomes of cultivated representatives of this family exceed 400 kbp and encode diverse functional repertoires including sphingolipid biosynthesis genes [44].

Although most orders contained primarily genomes that could be readily grouped into families, the *Pandoravirales* also included 15 singleton genomes out of the 37 total. This is potentially due to the lack of adequate genome sampling in this group, which would result in many distinct lineages represented by only individual genomes. If this is the case, more well-defined families will become evident as additional genomes are sequenced. Alternatively, the lack of clearly defined families could result from longer branches in this group that obfuscate the clustering of well-defined groups. The Medusavirus, which is included in this order, encodes a divergent PolB marker gene that is likely to be a replacement of the ancestral virus polymerase with a eukaryotic homolog [45]. Frequent gene transfers among the phylogenetic marker genes might be another explanation for the presence of many long branches in the *Pandoravirales* clade.

The *Pimascovirales* encompass 10 families including the *Iridoviridae* (PM_02)*, Marseilleviridae* (PM_05), and *Pithoviridae* (PM_07) and notably includes both *Pithovirus sibericum*, which has the largest viral capsid size currently known (1.5 μm [46]), as well as crustacean viruses in the family *Mininucleoviridae* (PM_10), which possess the smallest genomes recorded for any *Nucleocytoviricota* (67-71 kbp [47]). The *Mininucleoviridae* have highly degraded genomes that lack several phylogenetic marker genes. Although they can be classified within the *Pimascovirales* with high confidence, their relationship to other families is uncertain, and we therefore placed them in a polytomous node at the base of this order (Fig 2, see Methods). The uncharacterized family PM_01 contains the largest number of genomes (64) within this order, all of which are MAGs. The majority of these MAGs were derived from aquatic metagenomes, and some have been recovered in marine metatranscriptomes, suggesting they play an important but currently unknown role in marine systems. Overall, the repertoires of encoded proteins in the *Pimascovirales* were notably different from the *Imitervirales, Pandoravirales*, and *Algavirales*; while cytoskeleton components, nutrient transporters, light sensing genes, and central carbon metabolism components were prevalent in the latter three families, they were largely absent in the *Pimascovirales* (Fig 4b). Conversely, histone components appeared to be more prevalent in the latter order, whereas genes involved in translation and lipid metabolism were prevalent across most orders.

In addition to the families that fall within the established orders, we also identified two lineages that may represent novel orders or classes. The first is represented by two genomes that are basal-branching to the *Pimascovirales* (GVMAG-S-1041349-163 and GVMAG-M-3300025880-56, Fig 2). The placement of this lineage suggests that it might represent a new order that is sister lineage to the *Pimascovirales*. The second group is a set of 3 genomes that is basal-branching to the *Pokkesviricetes* class, which includes the orders *Chitovirales* and *Asfuvirales* (Fig 2). The basal-branching placement of this group suggests it might comprise a new class that is a sister group to the *Pokkesviricetes*. For both of these lineages, additional phylogenetic analyses with more genomic representatives will be necessary to confirm their evolutionary relationships to other *Nucleocytoviricota*.

## Discussion

Although only 6 families of *Nucleocytoviricota* have been established to date, recent cultivation-independent studies have revealed a vast diversity of these viruses in the environment, and their classification together with cultivated representatives has remained challenging. Here we present a unified taxonomic framework based on a benchmarked set of phylogenetic marker genes that establishes an hierarchical taxonomy of *Nucleocytoviricota*. This taxonomy encompasses 6 orders and 32 families, including one order and 26 families we propose here. Of the 32 families we demarcate, 30 fit into established orders, whereas two potentially represent new orders or even classes. Remarkably, the *Imitervirales* contain 11 families, including the *Mimiviridae*, underscoring the vast diversity of large viruses within this order. This framework substantially increases the total number of *Nucleocytoviricota* families, and we expect the number will continue to increase markedly as new genomes are incorporated. In particular, we identified 22 singleton genomes that likely represent additional families, the status of which will be clarified as more genomes become available.

We anticipate that the phylogenetic and taxonomic framework we develop here will be a useful community resource for several future lines of inquiry into the biology of *Nucleocytoviricota*. Firstly, the GVOGs are a large set of viral protein families constructed using many recently-produced *Nucleocytoviricota* MAGs, and they will likely be useful for the genome annotation and the examination of trends in gene content across viral groups. Secondly, the phylogenetic framework we present will facilitate work that delves into ancestral *Nucleocytoviricota* lineages, examines the timing and nature of gene acquisitions, and classifies newly discovered viruses. For example, giant viral genomes (> 500 kbp) evolved independently in multiple orders, and future studies that examine the similarities and differences in these genome expansion events will be important for pinpointing the driving forces of viral gigantism. Lastly, analysis of the environmental distribution of different taxonomic ranks of *Nucleocytoviricota* across Earth’s biomes will be an important direction for future work that reveals prominent biogeographic patterns and helps to clarify the ecological impact of these viruses.

## Methods

### *Nucleocytoviricota* genome set

We compiled a set of *Nucleocytoviricota* genomes that included metagenome-assembled genomes (MAGs) as well as genomes of cultured isolates. For this we first downloaded all MAGs available from several recent studies [9,15,20]. We also included all *Nucleocytoviricota* genomes available in NCBI RefSeq as of June 1st, 2020. Lastly, we also included several *Nucleocytoviricota* genomes from select publications that were not yet available in NCBI, such as the cPacV, ChoanoV, PoV01b, and AbALV viruses that have recently been described [14,33,48,49]. After compiling this set, we dereplicated the genomes, since the presence of highly similar or identical genomes is not necessary for broad-scale phylogenetic inference. For dereplication we compared all genomes against each other using MASH v. 2.0 [50] (“mash dist” parameters -k 16 and -s 300), and clustered genomes together using a single-linkage clustering, with all genomes with a MASH distance of ≥ 0.05 linked together. The MASH distance of 0.05 was chosen since it has been found to correspond to an average nucleotide identity (ANI) of 95% [50], which has furthermore been suggested to be a useful metric for distinguishing viral species-like clusters [51]. From each cluster, we chose the genome with the highest N50 contig length as the representative. We then decontaminated the genomes through analysis with ViralRecall v.2.0 [52] (-c parameter), with all contigs with negative scores removed on the grounds that they represent non-*Nucleocytoviricota* contamination or highly unusual gene composition that cannot be validated by our present knowledge of *Nucleocytoviricota* genomic content. We only considered contigs > 10 kbp given the inherent difficulty in eliminating contamination derived from short contigs. To ensure that we only used genomes that could be placed in a phylogeny, we then screened the genome set and retained only those with a PolB marker and three of the four markers A32, SFII, VLTF3, and MCP, consistent with our previous methodology. After this we arrived at a set of 1,380 genomes, including 1,253 MAGs and 127 complete genomes of cultivated viruses.

### GVOG construction

To construct GVOGs we first predicted proteins from all genomes using Prodigal v. 2.6.2. Proteins that did not have a recognizable start or stop codon at the ends of contigs were removed on the grounds that they may represent fragmented genes and obfuscate orthologous group predictions. We then calculated orthologous groups (OGs) using Proteinortho v. 6.06 [53] (parameters -e=1e-5 ╌identity=25 -p=blastp+ ╌selfblast ╌cov=50 -sim=0.80). We constructed Hidden Markov Models (HMMs) from proteins by aligning them with Clustal Omega v1.2.3 [54] (default parameters), trimming the alignment with trimAl v1.4.rev15 [55] (parameters -gt 0.1), and generating the HMM from the trimmed alignment with hmmbuild in HMMER v3.3 [56]. The goal of this analysis was to identify broad-level protein families, and we therefore sought to merge HMMs that bore similarity to each other and therefore derived from related protein families. For this we then compared the proteins in each OG to the HMM of every other OG (hmmsearch -E 1e-20 ╌domtblout option, hits retained only if 30% of the query protein aligned to the HMM). In cases where >50% of the proteins in one OG also had hits to the HMM of another OG, and vice versa, we then merged all of the proteins together and constructed a new merged HMM from the full set of proteins. The final set contained 8,863 HMMs, and we refer to these as the Giant Virus Orthologous Groups (GVOGs). To provide annotations for GVOGs we compared all of the proteins in each GVOG to the EggNOG 5.0 [57], Pfam [58], and NCVOG databases [59] (hmmsearch, -E 1e-3). For NCVOGs, we obtained protein sequences from the original NCVOG study and generated HMMs using the same methods we used for GVOGs. Annotations were assigned to a GVOG if >50% of the proteins used to make a GVOG had hits to the same HMM in one of these databases. Details regarding all GVOGs and their annotations can be found in Table S3.

### Benchmarking phylogenetic marker genes for *Nucleocytoviricota*

To identify phylogenetic markers for *Nucleocytoviricota* we cataloged GVOGs that were broadly represented in the 1,380 viral genomes that we used for benchmarking. We searched all proteins encoded in the genomes against the GVOG HMMs using hmmsearch (e-value cutoff 1e-10) and identified a set of 25 GVOGs that were found in >70% of the genomes in our set (hmmsearch, -E 1e-5). We constructed individual phylogenetic trees of these protein families to assess their individual evolutionary histories. For individual phylogenetic trees we calibrated bit score cutoffs so that poorly matching proteins would not be included. These cutoffs were equivalent to the 5th percentile score of all of the best protein matches for each genome. We then examined several features of these trees. Firstly, we only considered GVOGs present in all established families that would therefore be useful as universal or nearly-universal phylogenetic markers. Secondly, we examined each tree individually to assess the degree to which taxa from different orders clustered together in distinct monophyletic groups, which was taken as a signature of HGT. High levels of gene transfer would produce topologies incongruent with other marker genes and therefore compromise the reliability of a given marker when used on a concatenated alignment. For individual marker gene trees we aligned proteins from each GVOG using Clustal Omega, trimmed the alignment using trimAl (-gt 0.1 option), and constructed the phylogeny using IQ-TREE with ultrafast bootstraps calculated (-m TEST, -bb 1000, -wbt options).

We arrived at a set of 9 GVOGs that met the criteria described above and could potentially serve as robust phylogenetic markers (Table 1). We evaluated the phylogenetic strength of these markers individually using the recently-developed Tree Certainty (TC) and Internode Certainty (IC) metrics. These metrics are an improvement on the traditional bootstrap because they take into account the frequency of contrasting bipartitions, and can therefore be viewed as a measure of the phylogenetic strength of a gene [24,25]. We generated alignments using Clustal Omega, trimmed with TrimAl, and generated trees with IQ-TREE v1.6.9 [60] with ultrafast bootstraps [61] (parameters -wbt -bb 1000 -m LG+I+G4). We calculated TC and IC values in RaxML v8.2.12 (-f i option, ultrafast bootstraps used with the -z flag) [62]. We also evaluated the TC and IC values of trees generated from concatenated alignments. To construct concatenated alignments we used the python program ‘ncldv_markersearch.py’ that we developed for this purpose: https://github.com/faylward/ncldv_markersearch. For the 8 marker sets we evaluated we generated three replicate trees in IQ-TREE, and for comparisons between marker sets we used the trees that yielded the highest TC values.

For the final tree used for clade demarcation, we ran IQ-TREE five times using the parameters “-m LG+F+I+G4 -bb 1000 -wbt”, and we chose the resulting tree with the highest TC value for subsequent clade demarcation and RED calculation. Three genomes in the *Mininucleoviridae* family were included in the final tree but were not used for the benchmarking analysis because they have been shown to have highly degraded genomes that are not necessarily representative of *Nucleocytoviricota* more broadly [47]. Moreover, the MAG ERX555967.47 was found to have highly variable placement in different orders in different trees we analyzed, and we therefore did not include this genome in the final tree on the grounds that it represented a rogue taxa that may reduce overall tree quality [63].

### Family delineation and nomenclature

We calculated RED values in R using the get_reds function in the package “castor” [64]. As input we used a rooted tree derived from the 7-gene marker set described above. For the *Poxviridae, Asfarviridae, Iridoviridae, Phycodnaviridae, Marseilleviridae, Mininucleoviridae*, and *Mimiviridae* we retained existing nomenclature, and clades assigned these names based on the initially characterized viruses that were assigned to these families. For example, the *Phycodnaviridae* was assigned to AG_02 because the chloroviruses within this clade were the first-described members of this family, while the Prasinoviruses were assigned to a new family although they are commonly referred to as *Phycodnaviridae*. Similarly, *Mimiviridae* was assigned based on the placement of Acanthamoeba polyphaga mimivirus, *Iridoviridae* was assigned based on the placement of *Invertebrate iridescent virus 6*, *Asfarviridae* was assigned to the clade containing African swine fever virus (ASFV), and *Marseilleviridae* was assigned to the clade containing the Marseilleviruses. The treemap visualization was generated using the R package “treemap”.

## Data Availability

All data products described in this study are available on the Giant Virus Database: https://faylward.github.io/GVDB/

## Acknowledgments

We acknowledge the use of the Virginia Tech Advanced Research Computing Center for bioinformatic analyses performed in this study. This work was supported by grants from the Institute for Critical Technology and Applied Science and the NSF (IIBR-1918271) and a Simons Early Career Award in Marine Microbial Ecology and Evolution to F.O.A.. E.V.K. is supported by the funds of the Intramural Research Program of the National Institutes of Health (National Library of Medicine).

